# Socially stratified epigenetic profiles are associated with cognitive functioning in children and adolescents

**DOI:** 10.1101/2021.08.19.456979

**Authors:** L. Raffington, P.T. Tanksley, A. Sabhlok, L. Vinnik, T. Mallard, L.S. King, B. Goosby, K.P. Harden, E.M. Tucker-Drob

**Affiliations:** Department of Psychology, University of Texas at Austin, Austin, TX, USA.; Population Research Center, The University of Texas at Austin, Austin, TX, USA; Department of Psychiatry and Behavioral Sciences, Tulane University School of Medicine, New Orleans, LA, USA; Department of Sociology, University of Texas at Austin, Austin, TX 78712

**Keywords:** DNA-methylation, epigenetics, cognition, children, socioeconomic status, racism

## Abstract

Children’s cognitive functioning and educational performance are socially stratified. Social inequality, including classism and racism, may operate partly via epigenetic mechanisms that modulate neurocognitive development. Following preregistered analyses of data from 1,183 8-to 19-year-olds from the Texas Twin Project, we examined whether salivary DNA-methylation measures of inflammation (DNAm-CRP), cognitive functioning (Epigenetic-g), and pace of biological aging (DunedinPoAm) are socially stratified and associated with performance on tests of cognitive functions. We find that children growing up in more disadvantaged families and neighborhoods and children from marginalized racial/ethnic groups exhibit DNA-methylation profiles associated with higher chronic inflammation, lower cognitive functioning, and faster pace of biological aging. These salivary DNA-methylation profiles were associated with processing speed, general executive function, perceptual reasoning, verbal comprehension, reading, and math. Given that the DNA-methylation measures we examined were originally developed in adults, our results suggest that social inequalities may produce in children molecular signatures that, when observed in adults, are associated with chronic inflammation, advanced aging, and reduced cognitive function. Salivary DNA-methylation profiles might be useful as a surrogate endpoint in assessing the effectiveness of psychological and economic interventions that aim to reduce negative effects of childhood social inequality on lifespan development.

**Significance Statement:** Children’s cognitive functioning differs by dimensions of social inequality, such as class and race. Epigenetic mechanisms that regulate gene expression might be critically involved in the biological embedding of environmental privilege and adversity. We find that children growing up in more disadvantaged families and neighborhoods and from marginalized racial/ethnic groups exhibit higher chronic inflammation, lower cognitive functioning, and a faster pace of biological aging, as indicated by novel salivary DNA-methylation measures. These DNA-methylation measures of higher inflammation, lower cognitive functioning, and a faster pace of biological aging were, in turn, associated with performance on multiple cognitive tests. DNA-methylation measures might be useful as a surrogate endpoint in evaluation of programs to address the childhood social determinants of lifelong cognitive disparities.

## Introduction

Children’s cognitive function and educational performance are sensitive to environmental input, robustly predict their future social attainments and health (1), and consistently differ by major dimensions of social inequality, such as parental education, income, and race (2, 3). Socioeconomic and racial disparities in child cognitive development arise through various factors tied to classism and racism, including inequitable access to high-quality childcare, educational resources, healthcare, nutrition, and differences in exposure to toxicants, family stress, and neighborhood threat, among other factors (4, 5). For example, the social advantage of White identity, or White privilege, describes the generational legacy of social power experienced by White people through state-sanctioned social marginalization, which persistently shapes the disadvantaged context that young Black and Latinx youth face in the US. Due to the chronic nature of interpersonal and vicarious discrimination in their day-to day lives, indicators of socioeconomic disadvantage capture relevant but limited aspects of the effects of racism on child development (6).

Epigenetic mechanisms that regulate the expression of genes are hypothesized to be involved in the biological embedding of environmental privilege and disadvantage (7). For instance, a consistent finding from experimental manipulations of the social environment in nonhuman animals is that social adversity increases expression of genes linked to inflammation (8), which can modulate the continued development and function of the brain (7). Thus, epigenetic mechanisms actuated by classism and racism may, in part, contribute to social disparities in children’s cognitive function.

New advances of genome-wide technology and “omic” approaches have now quantified molecular signatures of a host of exposures, biological processes, and phenotypes that can be used to investigate the etiology of social disparities in life course development. For example, studies have identified patterns of DNA methylation across the epigenome in association with a peripheral proxy for systemic inflammation (9), multisystem biological aging processes (10), and psychological phenotypes (11). Results from such *discovery* studies can be used in *prediction* studies to construct epigenetic profiles in new samples that can then be examined in relation to a wide range of measured variables.

DNA-methylation discovery studies most commonly analyze methylation from blood or other tissues, but not methylation in salivary DNA, which comes from a mixture of buccal cells and leukocytes (https://ngdc.cncb.ac.cn/ewas/statistics). DNA methylation is a dynamic process and can be tissue-specific with, for example, different epigenetic signatures in brain, blood, and saliva (12). Yet, because DNA-methylation profiling using saliva is amenable at large scale in pediatric samples, this method offers distinct opportunities for large-scale epidemiological and longitudinal studies. Specifically, salivary DNA-methylation profiles may be useful for examining the etiology of social disparities in lifespan development. However, little work has been conducted to-date to examine whether salivary DNA-methylation measures are sensitive to social inequality (though see (13)) and associated with psychological development in children and adolescents.

Following preregistered analyses (https://osf.io/x978n/), we examined whether salivary DNA-methylation measures derived from discovery studies trained on inflammation, cognitive function, and the pace of biological aging are (a) stratified by major dimensions of social inequality and (b) associated with cognitive functions in children and adolescents. Three salivary DNA-methylation composite scores were of particular interest in the present study, because their blood-derived composites have been associated with cognitive function or they have been found to be sensitive to socioeconomic inequality. First, we examined DNA-methylation profiles of C-reactive protein (CRP; “DNAm-CRP”), which in blood samples have previously been found to be associated with cognitive functions in adults (14) and children (15). Second, we examined “Epigenetic-g” from a blood-based epigenome wide association study of general cognitive functions (*g*) in adults, which accounted for 3.4% and 4.5% of the variance in general cognitive functioning in two external adult cohorts using methylation from blood samples (11). Third, we examined “DunedinPoAm”, which was developed from analysis of rate of longitudinal change in organ system integrity occurring in middle-adulthood in a cohort of individuals who were all the same chronological age (16). We previously reported that socioeconomic disadvantage and Latinx compared to White identity is associated with faster pace of biological aging, as indicated by the DunedinPoAm, in a previous data freeze of Texas Twins salivary DNA-methylation data (N=600; (13)). In contrast, epigenetic clocks and the mortality predictor “GrimAge” were not sensitive to socioeconomic inequality and therefore not considered in analyses reported here (DNAm-CRP and Epigenetic-g were not previously examined). In the present study, we also examined whether genetic profiles of inflammation (*i.e.* polygenic scores of CRP, (17)) are associated with cognitive functions. Participants were 1183 (609 female) children and adolescents with at least one DNA-methylation sample from the population-representative Texas Twin Project, including 426 monozygotic and 757 dizygotic twins from 611 unique families, aged 8 to 19 years (mean age = 13.38 y, *SD* = 2.99 y).

## Results

The preregistration of our analysis plan can be found at https://osf.io/krgfs/.

### Salivary DNA-methylation profiles in children are reliably measured and show expected patterns of association with covariates

DNA-methylation profiles (*i.e.,* DNAm-CRP, Epigenetic-g, DunedinPoAm) measured from salivary DNA were approximately normally distributed (see **Table 1** for descriptive statistics before correction for the cell composition of saliva samples). Analyses of 15 technical replicates suggested moderate-to-good reliability of DNA-methylation profiles residualized for technical artifacts and cell composition (ICC for DNAm-CRP = 0.73, epigenetic-g = 0.80, DunedinPoAm = 0.84). Biometric models using the twin family structure, where the similarity between twins due to both additive genetic factors (*A*) and environmental factors shared by twins living in the same home (*C*) represents a lower bound estimate of reliability, also suggested good reliability of DNA-methylation profiles (*A*+*C* variation for DNAm-CRP= 60.7%, Epigenetic-g = 55.3%, DunedinPoAm = 54.2%, accounting for age and gender).

**Table 1.** Descriptive statistics.

| Sample | *n* | *M* | *SD* |
| --- | --- | --- | --- |
| DNAm-CRP<sup>a</sup> | 1365 | 0.03 | 0.02 |
| Epigenetic-g<sup>a</sup> | 1365 | -0.02 | 0.20 |
| DunedinPoAm<sup>a</sup> | 1365 | 1.02 | 0.06 |
| PGS-CRP<sup>b</sup> | 654 | 0 | 1 |
| Processing speed | 869 | 0 | 1 |
| General executive functions | 869 | 0 | 1 |
| Perceptual reasoning | 1328 | 104.18 | 14.48 |
| Verbal comprehension | 1329 | 105.61 | 14.02 |
| Reading | 625 | 30.97 | 4.9 |
| Math | 619 | 21.24 | 6.48 |
| Family-level socioeconomic disadvantage | 993 | -0.02 | 0.96 |
| Neighborhood-level socioeconomic disadvantage | 1218 | -0.02 | 1 |
| Neighborhood opportunity | 950 | 0.31 | 0.62 |
| Body mass index | 1364 | 0.4 | 1.34 |
| Pubertal development | 1325 | 2.61 | 0.92 |
| Tobacco use (yes/no) | 58/631 | — | — |
| DNAm-smoke | 1365 | -13.62 | 1.84 |

<sup>a</sup> After exclusion of participants based on DNA-methylation preprocessing (n=44), excluding technical replicates (n=15), and including repeated samples (n= 182). Means of raw scores before residualizing for cell composition, array, slide, and batch. Scores were standardized (mean = 0, SD = 1) for analyses.
<sup>b</sup> PGS-CRP only computed for individuals solely of recent European ancestries.

Higher DNAm-CRP was strongly correlated with higher DunedinPoAm (*r*=0.89 [95% CI= 0.81, 0.96], *p*<0.001, accounting for age and gender) and moderately correlated with lower Epigenetic-g (*r*=-0.31 [-0.42, -0.19], *p*<0.001). This result is unsurprising as CRP levels were one of the 18 biomarkers that the DunedinPoAm measure was trained on (16). Lower Epigenetic-g was weakly correlated with higher DunedinPoAm (*r*=-0.17 [-0.29, -0.04], *p*=0.011).

Older children had higher DNAm-CRP (*r* =0.35 [0.26, 0.44], *p*<0.001), Epigenetic-g (*r*=0.64 [0.56, 0.72], *p*<0.001), and DunedinPoAm profiles (*r*=0.13 [0.02, 0.23], *p*=0.018). Boys had lower DNAm-CRP (*d* =-0.26 [-0.34, -0.18], *p*<0.001) and DunedinPoAm (*d* =-0.18 [-0.27 - 0.10], *p*<0.001), but not Epigenetic-g profiles (*d* =0.06 [-0.02, 0.14], *p*=0.143). All models included age, gender, and an age by gender interaction as covariates.

### Salivary DNA-methylation profiles are socially stratified in children

Salivary DNA-methylation profiles in children were socially stratified. Children from socioeconomically disadvantaged families, socioeconomically disadvantaged neighborhoods, neighborhoods with less intergenerational economic mobility (*i.e.,* neighborhood opportunity), and children reporting Latinx-only or Black+ identity relative to White-only identity exhibited DNA-methylation profiles associated with higher chronic inflammation, a faster pace of biological aging, and lower cognitive functioning (see **Figure 1** and **Supplemental Table S1**). Children reporting Black+ and Latinx-only identities lived in the most socioeconomically disadvantaged families and neighborhoods compared to children reporting White-only identity (see **Figure 2** and **Supplemental Table S2**). Family-level socioeconomic disadvantage accounted for racial/ethnic disparities in DNAm-CRP, but not Epigenetic-g. Family-level socioeconomic disadvantage accounted for the difference in DunedinPoAm between Black+, but not Latinx-only, relative to White-only identity (see **Supplemental Table S1**). See **Supplemental Table S3** for effect size estimates between socioeconomic inequality and DNA-methylation profiles reported separately for each racial/ethnic group (this analysis was not preregistered).

**Figure 1.**
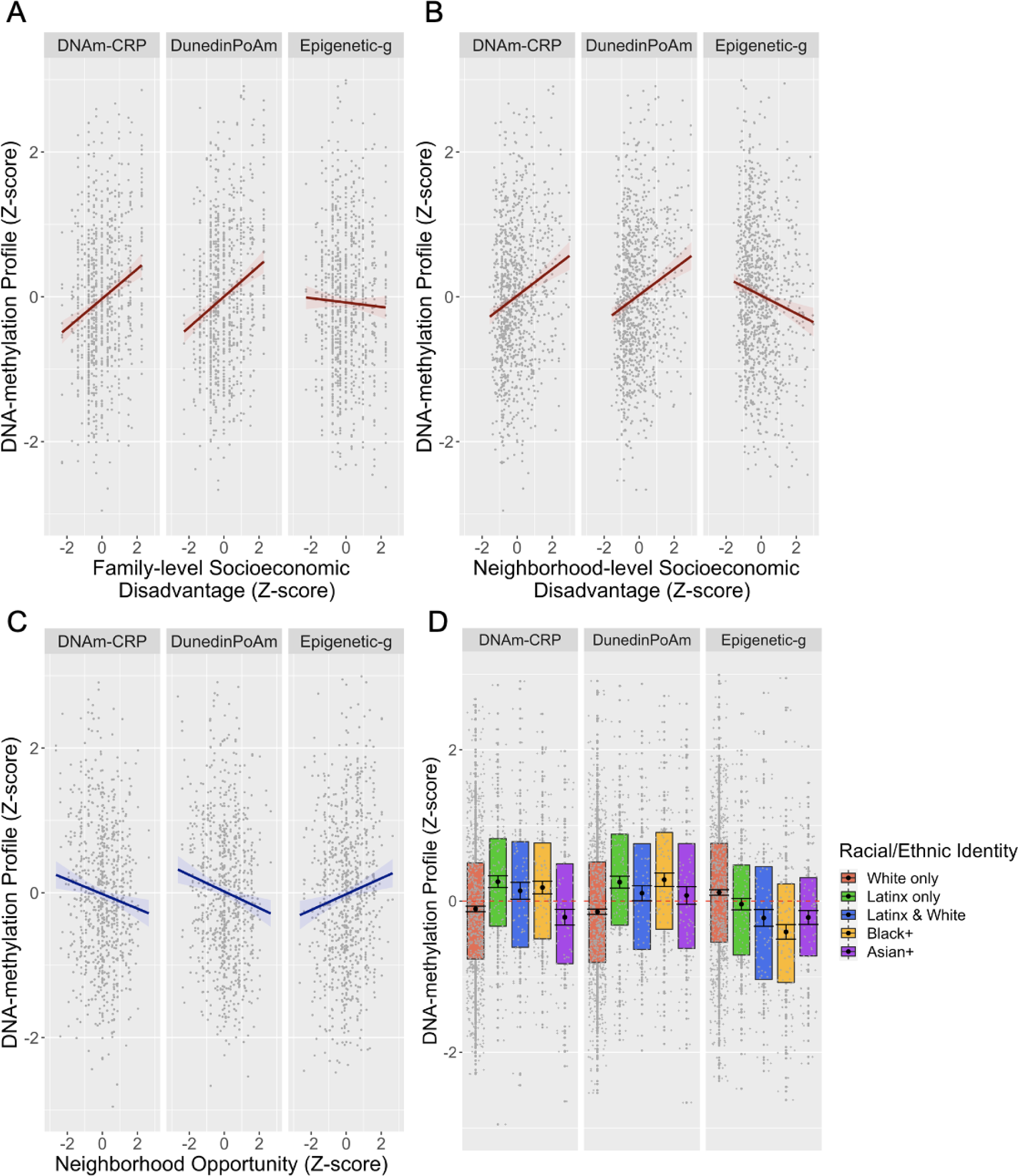
Associations between A) family-level socioeconomic disadvantage, B) neighborhood-level socioeconomic disadvantage, C) neighborhood opportunity (*i.e.,* intergenerational economic mobility), and D) self-identified racial/ethnic identity with three DNA-methylation profiles (DNAm-CRP, DunedinPoAm, and Epigenetic-g) in children and adolescents. DNA-methylation profiles and socioeconomic disadvantage values are in standard deviation units. Higher DNAm-CRP values indicate a methylation profile associated with higher chronic inflammation. Higher DunedinPoAm values indicate a methylation profile associated with faster biological aging. Higher Epigenetic-g values indicate a methylation profile associated with higher cognitive functioning. The racial/ethnic identity boxplots display group DNA-methylation differences in the mean (black circle), standard errors of the mean (error bars), the first and third quartiles (lower and upper hinges), and the mean across groups (red dashed line). Participants self-identified as White only (62%), Latinx only (12.2%), Latinx and White (8.1%), Black and potentially another race/ethnicity (10%), Asian and potentially another race/ethnicity but not Latinx or Black (7.5%), and Indigenous American, Pacific Islander or other, but not Latinx, Black, or Asian (0.6%, not shown due to small sample size). See **Supplemental Table S1** for standardized regression coefficients with and without covariate controls for body mas index, puberty, and socioeconomic inequality.

**Figure 2.**
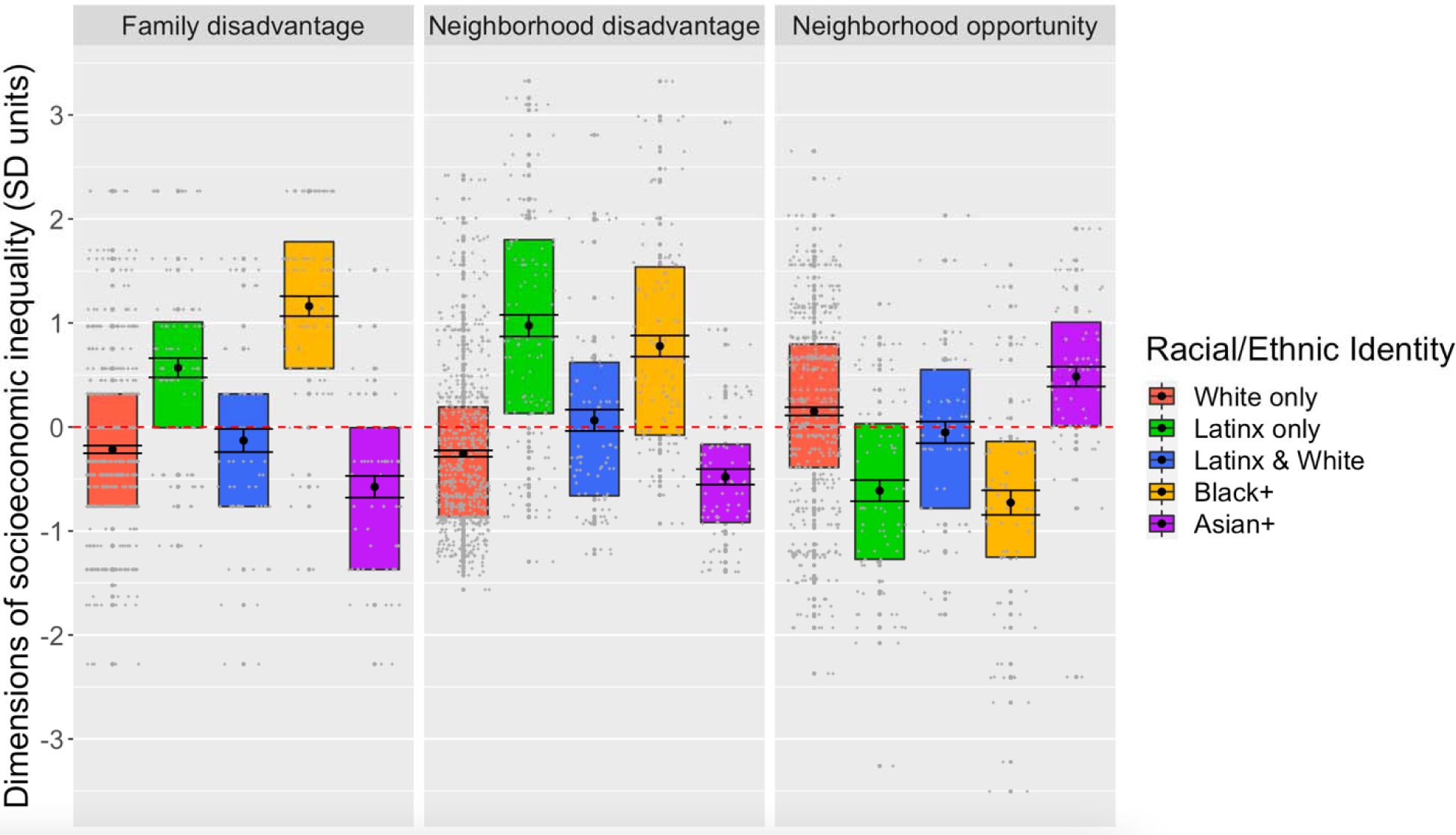
Associations between racial/ethnic identity and dimensions of socioeconomic inequality. Socioeconomic disadvantage and opportunity values are Z-scores. The racial/ethnic identity boxplots display group DNA-methylation differences in the mean (black circle), standard errors of the mean (error bars), the first and third quartiles (lower and upper hinges), and the mean across groups (red dashed line). Participants self-identified as White only (62%), Latinx only (12.2%), Latinx and White (8.1%), Black and potentially another race/ethnicity (10%), Asian and potentially another race/ethnicity but not Latinx or Black (7.5%), and Indigenous American, Pacific Islander or other, but not Latinx, Black, or Asian (0.6%, not shown due to small sample size). See **Supplemental Table S2** for standardized regression coefficients.

Comparing White-only identifying children to all other groups (this comparison was not preregistered) indicated that the advantage, or privilege, of White identity compared to other racial/ethnic categories was evident in all three DNA-methylation profiles (DNAm-CRP *r* = - 0.14 [-0.22, -0.06], *p*<0.001; Epigenetic-g *r*= 0.23 [0.16, 0.31], *p*< 0.001; DunedinPoAm *r*=-0.25 [-0.34, -0.16], *p*<0.001). White identity remained evident in Epigenetic-g (*r*=0.21 [0.13, 0.29], *p*<0.001) and DunedinPoAm (*r*=-0.19 [-0.29, -0.09], *p*<0.001), but not DNAm-CRP (*r*=-0.08 [-0.17, 0.01], *p*=0.067), after accounting for the lower rates of family-level disadvantage experienced by White children (*r*=-0.29 [-0.38, -0.19], *p*<0.001). Effects of White identity were reduced but also still remained evident in Epigenetic-g (*r*=0.17 [0.09–0.25], *p*<0.001) and DunedinPoAm (*r* =-0.16, [-0.26, -0.05], *p*=0.003), but not DNAm-CRP (*r*=-0.04 [-0.13, 0.05], *p*=0.373), after accounting for both the lower rates of family-level (*r*=-0.30 [-0.39, -0.21], *p*<0.001) and neighborhood-level socioeconomic disadvantage (*r*=-0.34 [-0.42, -0.26], *p*<0.001) experienced by White children.

We next examined the role of body mass index (BMI), pubertal stage, and DNA-methylation profiles related to smoking (DNAm-smoke) in associations of social inequality and DNA-methylation profiles of interest. Socioeconomic and racial/ethnic inequalities in DNA-methylation largely remained after including these covariates, with the exception that correlations with DNAm-CRP were largely accounted for by including BMI (see **Supplemental Table S1**).

### Salivary DNA-methylation profiles are associated with cognitive functions

Salivary DNA-methylation profiles were associated with performance on multiple in-laboratory tests of cognitive functioning: Higher DNAm-CRP was associated with worse performance on tests of processing speed, general executive function, perceptual reasoning, and verbal comprehension. Lower Epigenetic-g was associated with lower scores on tests of perceptual reasoning, verbal comprehension, reading, and math. Finally, faster pace of biological aging was associated with lower scores on tests of verbal comprehension and perceptual reasoning (see **Figure 3** and **Supplemental Table S4**). Notably, the largest effect size was observed for Epigenetic-g and math, where Epigenetic-g explained R^2^=11.1% of the variation in math performance. See **Supplemental Table S5** for effect size estimates between DNA-methylation profiles and cognition reported separately for each racial/ethnic group (this analysis was not preregistered).

**Figure 3.**
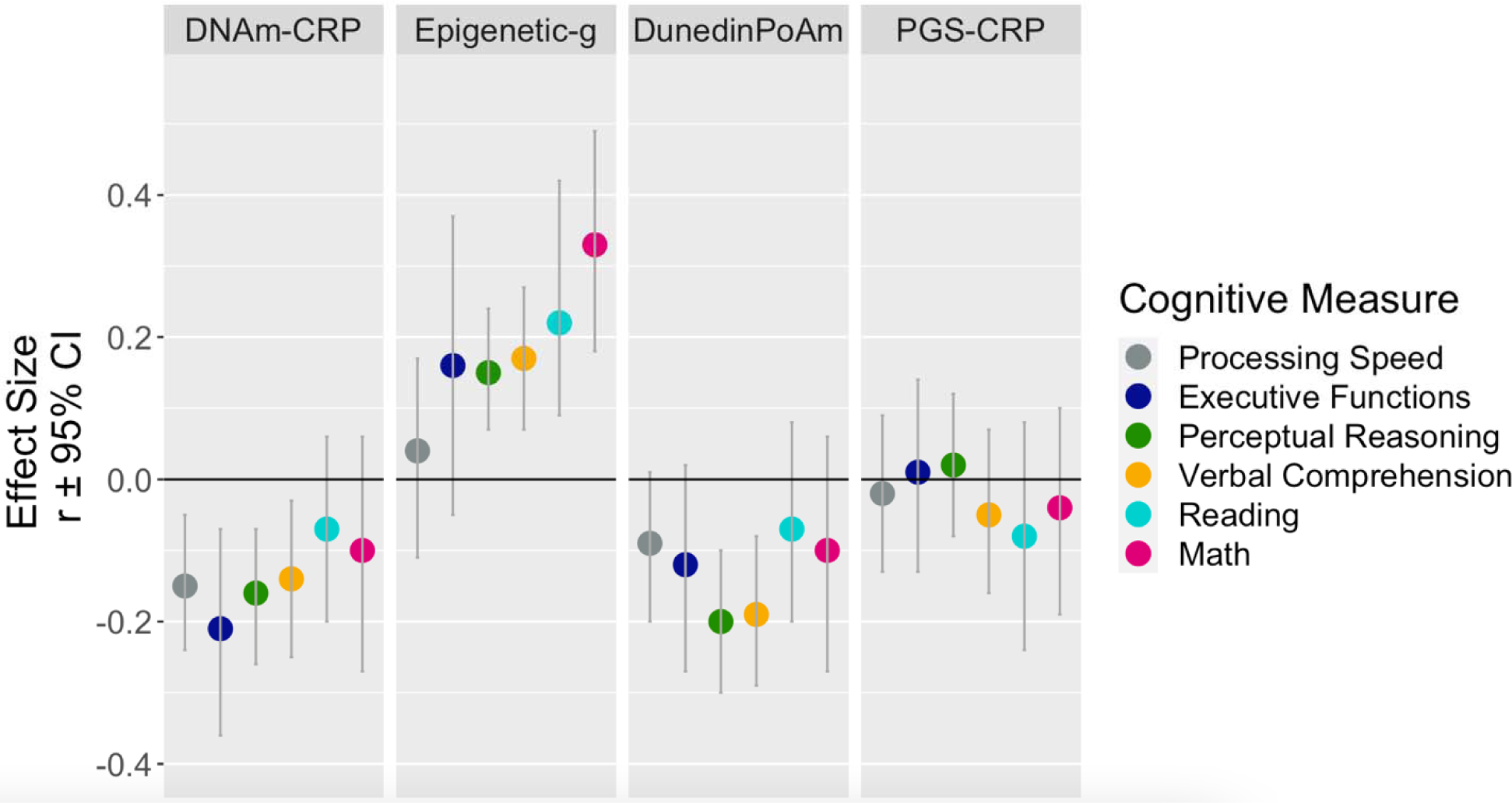
Associations between three DNA-methylation profiles (DNAm-CRP, Epigenetic-g, and DunedinPoAm) and a polygenic score of inflammation (PGS-CRP) with six measure of cognitive functioning in children and adolescents. The plot depicts the standardized regression coefficients (*r*) and 95% confidence intervals (CIs) calculated by regressing cognitive functions on DNA-methylation measures and PGS-CRP, separately. PGS analyses were restricted to participants solely of recent European ancestries as indicated by genetic ancestry PCs that are comparable to the GWAS discovery sample. All models included covariate adjustment for child’s age and gender, and technical covariates. Higher cognitive values indicat higher task performance. Higher DNAm-CRP and PGS-CRP values indicate a methylation profile and genetic profile of higher chronic inflammation, respectively. Higher Epigenetic-g values indicate a methylation profile associated with higher cognitive functioning. Higher DunedinPoAm values indicate a methylation profile of faster biological aging. See **Supplemental Table S4** for standardized regression coefficients with and without covariate controls for body mass index, puberty, and socioeconomic inequality.

As all three DNA-methylation profiles were associated with perceptual reasoning and verbal comprehension, we performed commonality analyses to examine the proportion of overlapping and unique variation explained. DNAm-CRP and DunedinPoAm explained largely overlapping variation in perceptual reasoning (DNAm-CRP alone: 2.6%, DunedinPoAm alone: 4%, combined: 3.8%) and verbal comprehension (DNAm-CRP alone: 2%, DunedinPoAm alone: 3.4%, combined: 3.8%). Whereas Epigenetic-g explained unique variation in perceptual reasoning (Epigenetic-g alone: 2.4%) and verbal comprehension (Epigenetic-g alone: 2.9%) relative to both DNAm-CRP (perceptual reasoning combined: 5.9%, verbal comprehension combined: 5.5%) and DunedinPoAm (perceptual reasoning combined: 6.5%, verbal comprehension combined: 6.3%).

We next examined the role of BMI, puberty, DNAm-smoke, and family-level disadvantage in associations of DNA-methylation measures with cognitive test performance. Associations were largely unaffected by controlling for BMI, puberty, and DNAm-smoke, with the exception that associations of DNAm-CRP with cognition were mostly accounted for by controlling for BMI. Associations of DNAm-CRP and DunedinPoAm with cognition were largely accounted for by controlling for family-level disadvantage, with the exception of perceptual reasoning. In contrast, associations of Epigenetic-g with cognitive test performance were unaffected by controlling for family-level disadvantage (**Supplemental Table S4**).

We assessed potential effects of differing sample sizes of cognitive measures (**Table 1**) on effect size estimates (this analysis was not preregistered). Effect size estimates based on models using listwise deletion were largely similar to reported results, suggesting that differing sample sizes across measures did not substantially affect effect sizes (see **Supplemental Figure 1**).

We further examined the extent to which DNA-methylation associations with cognition are robust to complete genetic and family-level environmental control in a bivariate biometric model that used the twin family structure of the Texas Twin Project. Consistent with the hypothesis that DNA-methylation associations with cognitive function represents (partially unmeasured) effects of family-level stratification, we found no evidence to suggest that identical twins who differ from their co-twins in DNA-methylation show corresponding differences in their cognitive functioning (see **Supplemental Table S6**).

### Genetic profiles of CRP are not associated with cognitive functions

PGS-CRP analyses were restricted to participants who had European genetic ancestries as indicated by principal components of genetic ancestry that were comparable to the GWAS discovery sample. PGS-CRP were not associated with measures of cognitive functioning (see **Figure 1** and **Supplemental Table S4**). PGS-CRP did not account for differences in cognitive function between dizygotic twins (see **Supplemental Figure S1**). Because of the sample size of dizygotic twin pairs (N=364) we preregistered the previous analysis as primarily exploratory.

## Discussion

We analyzed salivary DNA-methylation data from 1183 children and adolescents participating in the Texas Twin Project to examine whether salivary DNA-methylation measures of inflammation, cognitive function, and the pace of aging are (a) stratified by major dimensions of social inequality and (b) associated with performance on test of cognitive functions in childhood. We found that children and adolescents growing up in more socioeconomically disadvantaged families and neighborhoods and children from marginalized racial/ethnic groups compared to their more privileged peers exhibit DNA-methylation profiles associated with higher chronic inflammation, lower cognitive functioning, and a faster pace of biological aging. Moreover, these socially stratified DNA-methylation profiles were related to scores on multiple in-laboratory cognitive tests, including tests of processing speed, general executive function, perceptual reasoning, verbal comprehension, reading, and math. Associations of DNA-methylation measures of inflammation and the pace of aging with cognition were largely accounted for by controlling for family-level socioeconomic disadvantage.

Given that the DNA-methylation measures we examined were originally developed in adults, our results suggest that social inequalities may produce in children molecular signatures that, when observed in adults, are associated with chronic inflammation, advanced aging, and reduced cognitive function. Our findings indicate that salivary DNA-methylation measures, originally validated in adult blood samples, may be useful for indexing social inequality and risk for disparities in cognitive function in childhood and adolescence, sensitive developmental periods in which cognitive functions are susceptible to environmental inputs. Our cross-sectional, observational design did not allow us to examine whether policy changes mitigating socioeconomic inequality (*e.g.,* increases in minimum wage, child tax credits) and structural racism (*e.g.,* eliminating the legacy of redlining, police reforms) affects children’s DNA-methylation profiles. Such investigations remain high-priority areas for future research. Salivary DNA-methylation measures may be useful as surrogate endpoints for assessing the effectiveness of programs and policies that aim to reduce effects of childhood social inequality on lifespan development.

Our analysis of racial/ethnic group differences found that children reporting Latinx-only or Black+ relative to White-only social identity exhibited higher chronic inflammation, faster pace of biological aging, and lower cognitive functioning, as indicated by DNA-methylation measures. Children reporting Black+ or Latinx-only identity lived, on average, in substantially more socioeconomically disadvantaged families and neighborhoods compared to children reporting White-only identity. Socioeconomic disadvantage statistically accounted for some, but not all, of the differences between racial/ethnic groups in DNA-methylation profiles. For example, the social advantage of White identity, or White privilege, remained evident in DNA-methylation profiles after accounting for the lower rates of both family-level and neighborhood-level disadvantage experienced by White families. Thus, our findings are consistent with observations that racial and ethnic disparities leave biological traces in the first two decades of life and reflect multiple dimensions of social inequality (4, 5). Family and neighbourhood indicators of socioeconomic disadvantage, privilege, and intergenerational mobility capture relevant but limited aspects of the effects of racism on child development (6). Additional factors that are often neglected, such as the impact of race-based discrimination in education and healthcare systems and chronic exposure to interpersonal and vicarious discrimination in daily life, may explain further variance in the effects of racial/ethnic marginalization (18–20). Given that research in developmental psychology investigating the role of race, including racial disparities in adversity, is rare, scientific understanding of how racism manifests in children’s lives and affects their development remains limited (21).

We found that salivary DNA-methylation profiles were associated with several measures of cognitive functioning with non-negligible effect sizes. After correcting for multiple comparisons, DNA-methylation profiles of higher inflammation were associated with lower in-laboratory processing speed, general executive function, perceptual reasoning, and verbal comprehension. Lower Epigenetic-g was associated with lower perceptual reasoning, verbal comprehension, reading, and math performance. Faster pace of biological aging was correlated with lower verbal comprehension and perceptual reasoning. Notably, Epigenetic-g explained 11.1% of the variation in math performance.

DNA methylation is a dynamic process and can be tissue specific with, for example, different epigenetic signatures in brain, blood, and saliva. Whereas we measured methylation in salivary DNA, the original estimates on which our profiles were based were estimated from DNA methylation in blood. Recent research suggests that salivary DNA-methylation collected with Oragene kits (as was done here) in children is particularly enriched for immune cells rather than epithelial cells (22). It may therefore be particularly sensitive to inflammatory processes, which contribute to DNA-methylation profiles of inflammation (DNAm-CRP) and pace of aging (DunedinPoAm). In contrast, genetic profiles related to inflammation (*i.e.,* polygenic scores of CRP) were not associated with cognitive functioning. Our findings linking DNAm-CRP with DunedinPoAm and cognitive functioning are in line with experimental animal studies reporting that social adversity increases expression of genes linked to inflammation, which may be critically involved in multi-system aging processes (*i.e.*, “inflammaging”) and can modulate the development of the brain (7, 8). Yet, the measurements we studied are molecular derivatives of unobserved inflammatory processes, not direct observations of chronic inflammation. Accordingly, this type of omics research is not well-suited to identifying precise biological processes.

In conclusion, our findings suggest that salivary DNA-methylation profiles are promising candidate biomarkers of major dimensions of social inequality experienced in real-time during childhood. Because saliva can easily be collected in large-scale pediatric epidemiological studies, salivary DNA-methylation profiles might be useful as surrogate endpoints in evaluation of ontogenetic theories and social programs that address the childhood social determinants of lifelong cognitive disparities.

## Method

### Sample

#### The Texas Twins Project

The Texas Twin Project is an ongoing longitudinal study that includes the collection of saliva samples for DNA and DNA-methylation extraction since 2012 (23). Participants in the current study were 1213 (622 female) children and adolescents, including 433 monozygotic and 780 dizygotic twins (see zygosity measure) from 617 unique families, aged 8 to 19 years (*M* = 13.66, *SD* = 3.06) that had at least one DNA-methylation sample. 195 participants contributed two DNA-methylation samples (time between repeated samples: *M* = 22 months, *SD* = 6.5, range 3 – 38 months) and 16 samples were assayed in duplicate for reliability analyses (total methylation sample n = 1424). Participants self-identified as White only (62%), Latinx only (12.2%), Latinx and White (8.1%), Black and potentially another race/ethnicity (10%), Asian and potentially another race/ethnicity but not Latinx or Black (7.5%), and Indigenous American, Pacific Islander or other, but not Latinx, Black, or Asian (0.6%). The University of Texas Institutional Review board granted ethical approval. Please see **Table 1** for descriptive statistics, including sample sizes for each measure after exclusion based on DNA-methylation preprocessing.

### Measures

#### DNA-methylation

##### DNA-methylation preprocessing

Saliva samples were collected during a laboratory visit using Oragene kits (DNA Genotek, Ottawa, ON, Canada). DNA extraction and methylation profiling was conducted by Edinburgh Clinical Research Facility (UK). The Infinium MethylationEPIC BeadChip kit (Illumina, Inc., San Diego, CA) was used to assess methylation levels at 850,000 methylation sites. DNA methylation preprocessing was primarily conducted with the ‘minfi’ package in R (24). Within-array normalization was performed to address array background correction, red/green dye bias, and probe type I/II correction, and it has been noted that at least part of the probe type bias is a combination of the first two factors (Dedeurwaerder et al., 2014). Noob preprocessing as implemented by minfi’s “preprocessNoob” (25) is a background correction and dye-bias equalization method that has similar within-array normalization effects on the data as probe type correction methods such as BMIQ (Teschendorff et al., 2013).

In line with our preregistered preprocessing plan, CpG probes with detection p > 0.01 and fewer than 3 beads in more than 1% of the samples and probes in cross-reactive regions were excluded (26). None of these failed probes overlapped with the probes used for DNA-methylation scores. 44 samples were excluded because (1) they showed low intensity probes as indicated by the log of average methylation <9 and their detection p was > 0.01 in >10% of their probes, (2) their self-reported and methylation-estimated sex mismatch, and/or (3) their self-reported and DNA-estimated sex mismatch. Cell composition of immune and epithelial cell types (*i.e.,* CD4+ T-cell, natural killer cells, neutrophils, eosinophils, B cells, monocytes, CD8+ T-cell, and granulocytes) were estimated using a newly developed child saliva reference panel implemented in the R package “BeadSorted.Saliva.EPIC” within “ewastools” (22). Surrogate variable analysis was used to correct methylation values for batch effects using the “combat” function in the SVA package (27).

#### DNA-methylation scores

##### DNAm-CRP

DNAm-CRP was computed on the basis of an epigenome-wide association study of CRP (9). Using the summary statistics of the associations between CpG sites and adult CRP, we created one methylation score per person by summing the product of the weight and the individual beta estimate for each individual at each of the 218 CpG sites significantly associated (*p□*<*□*1.15*□*×*□*10^−7^) with CRP.

##### DunedinPoAm

DunedinPoAm was developed from DNA-methylation analysis of Pace of Aging in the Dunedin Study birth cohort. Pace of Aging is a composite phenotype derived from analysis of longitudinal change in 18 biomarkers of organ-system integrity measured when Dunedin Study members were all 26, 32, and 38 years of age (28). Elastic-net regression machine learning analysis was used to fit Pace of Aging to Illumina 450k DNA-methylation data generated from blood samples collected when participants were aged 38 years. The elastic net regression produced a 46-CpG algorithm. Increments of DunedinPoAm correspond to “years” of physiological change occurring per 12-months of chronological time. The Dunedin Study mean was 1, *i.e.* the typical pace of aging among 38-year-olds in that birth cohort. Thus, 0.01 increment of DunedinPoAm corresponds to a percentage point increase or decrease in an individual’s pace of aging relative to the Dunedin birth cohort at midlife. DunedinPoAm was calculated based on the published algorithm (16) using code available at https://github.com/danbelsky/DunedinPoAm38.

##### Epigenetic-g

Salivary “epigenetic-g” was computed on the basis of a blood-based epigenome wide association study of general cognitive functions (*g*) in adults (11). We calculated epigenetic-g based on the algorithm available at https://gitlab.com/danielmccartney/ewas_of_cognitive_function. Prior to computation, methylation values were scaled within each CpG site (mean = 0, SD = 1). All DNA-methylation scores were residualized for array, slide, batch, cell composition and then standardized to ease interpretation.

#### Genetics

##### Genotyping and imputation

DNA samples were genotyped at the University of Edinburgh using the Illumina Infinium PsychArray, which assays ∼590,000 single nucleotide polymorphisms (SNPs), insertions-deletions (indels), copy number variants (CNVs), structural variants, and germline variants across the genome. Genetic data was subjected to quality control procedures recommended for chip-based genomic data (29, 30). Briefly, samples were excluded on the basis of poor call rate (< 98%) or inconsistent self-reported and biological sex, while variants were excluded if missingness exceeded 2%. As further variant-level filtering has been shown to have a detrimental effect on imputation quality (31), quality control thresholds for minor allele frequency (MAF) and Hardy–Weinberg equilibrium (HWE) were applied after phasing and imputation.

Untyped markers were imputed on the Michigan Imputation Server (https://imputationserver.sph.umich.edu). Specifically, genotypes were phased and imputed with Eagle v2.4 and Minimac4 (v1.5.7), respectively, while using the 1000 Genomes Phase 3 v5 reference panel (32). To ensure that only high-quality typed and imputed markers were used for analysis, variants were excluded if they had a MAF < 1e-3, a HWE *p*-value < 1e-6, or an imputation quality score < .90. These procedures produced a final set of 4,703,309 genetic markers to be used in analyses.

##### DNA preprocessing

DNA samples were genotyped at the University of Edinburgh using the Illumina Infinium PsychArray, which assays ∼590,000 single nucleotide polymorphisms (SNPs), insertions-deletions (indels), copy number variants (CNVs), structural variants, and germline variants across the genome. Genotypes will be subjected to quality control procedures recommended for chip-based genomic data (29, 30). Briefly, samples will be excluded due to poor call rate (< 98%) and inconsistent self-reported and biological sex. Variants will be excluded if more than 2% of data is missing. Untyped variants will be imputed on the Michigan Imputation Server (https://imputationserver.sph.umich.edu). As part of this process, genotypes will be phased with Eagle v2.4 and imputed with Minimac4 (v1.5.7), using the 1K Genomes Phase 3 v5 panel as a reference panel (32). Thresholds for minor allele frequency (MAF < 1e-3) and Hardy-Weinberg Equilibrium (HWE p-value < 1e-6) will be applied after phasing and imputation, as variant-level filtering has been shown to have a detrimental effect on imputation quality (31). Finally, imputed genotypes will be excluded if they suffer from poor imputation quality (INFO score < .90).

##### Polygenic scores

PGS-CRP was computed in two steps. First, GWAS summary statistics were adjusted for linkage disequilibrium, or LD (i.e., correlation structures in the genome that capture population stratification). The preregistered analysis plan proposed using SBayesR (33) for LD-adjustment. However, as the GWAS summary statistics used to compute PGS-CRP did not meet the data requirements of SBayesR (e.g., effect allele frequency, per SNP sample size), we elected to use PRScs for LD-adjustment instead. PRScs is a program that uses Bayesian regression to infer posterior SNP effects using continuous shrinkage priors. PRScs has been shown to improve prediction accuracy of PGSs over other widely used PGS approaches (34). PRScs requires GWAS summary statistics and an external reference panel of the same ancestry as the GWAS. For the summary statistics, we used publicly available data from a GWAS of CRP in 204,402 individuals solely of European ancestry (17). For the reference panel, we used the 1000 Genomes Project (32) European reference panel (phase 3 v5; provided with the software) that was restricted to HapMap3 SNPs (35).

Second, we used PLINK v2 (36) to apply the LD-adjusted SNP effects from PRScs in the Texas Twins Project sample. The resulting PGS-CRP is described by the following equation:

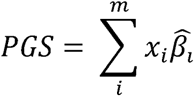

where *m* is the number of SNPs, 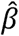 is the estimated effect of the *i*th SNP and *x*, coded as 0, 1 or 2, is the number of effect alleles of the th SNP. All PGS analyses were restricted to individuals solely of European ancestries in order to reduce the risk of spurious findings due to population stratification. PGS-CRP was residualized for the top five genetic principal components and genotyping batch and then standardized using Z-scores (M=1; SD=0).

#### Cognitive function

##### Processing speed

Three tasks were used to construct a latent measure of processing speed that were available in participants in grades three through eight: Symbol Search (Wechsler, 2003) Pattern Comparison, and Letter Comparison (Salthouse & Babcock, 1991). Each task assessed how quickly and accurately participants identified similarities between symbols, patterns, or letters.

##### Executive functions

The current study included 15 tasks assessing 4 EF domains that were available in participants in grades three through eight: inhibition, switching, working memory, and updating. Tasks were administered orally, on the computer, or on paper. Inhibition was assessed with four tasks: Animal Stroop (39), Mickey (40), and Stop Signal. The study originally used an auditory Stop Signal task (41), which was replaced with a visual Stop Signal task (42) after the third year of data collection to accommodate the needs of administering EF tasks in the MRI scanner. Switching was assessed using four tasks: Trail Making (Salthouse, 2011), Local-Global (44), Plus-Minus (44), and a computerized Cognitive Flexibility task (45). Cognitive Flexibility replaced the Plus-Minus task, again to accommodate MRI task administration after the third year of data collection. Working memory was assessed using three tasks: Symmetry Span (46), Digit Span Backward (37), and Listening Recall (47). These tasks tap spatial, verbal, and auditory working memory, respectively. Updating was assessed with four tasks: Keeping Track (44), Running Memory for Letters (48), 2-Back task (49), and, as a replacement to the 2-Back task after the third year of data collection, a 1- and 2-back task (49). More comprehensive task descriptions can be found in Engelhardt et al. (50).

Previous research in this sample (50, 51) demonstrated that variation in EF is best captured by a hierarchical factor model, with individual EF tasks loading onto one of four latent factors representing each EF domain and each of these loading onto a common EF factor. This same hierarchical model was adopted in all the analyses presented in the current research to examine general EF.

##### Verbal comprehension and Perceptual reasoning

We administered the Wechsler Abbreviated Scale of Intelligence (WASI-II; Wechsler, 2011) to all participants to assess perceptual reasoning, also called non-verbal fluid intelligence, and verbal comprehension, also called verbal crystallized intelligence. Perceptual reasoning is the sum of the age-normed t-scores on the Block Design and Matrix Reasoning subtests. Verbal comprehension is the sum of the age-normed t-scores on the Vocabulary and Similarities subtests.

##### Math and reading

To assess more specific reading comprehension and mathematics skills, participants in grades three through eight completed the Passage Comprehension and Calculation subtests, respectively, of the Woodcock-Johnson III Tests of Academic Achievement (53). The dependent variable for the reading and math subtests is total number of items correct.

#### Socioeconomic context

##### Family-level socioeconomic disadvantage

The family-level measure was computed from parent reports of household income, parental education, occupation, history of financial problems, food insecurity (based on the US Household Food Security Survey Module (54)), father absence, residential instability (changes in home address), and family receipt of public assistance. These were aggregated to form a composite measure of household-level cumulative socioeconomic disadvantage described in (2), and coded such that higher scores reflect greater disadvantage.

##### Neighborhood-level socioeconomic disadvantage

The neighbourhood-level measure was composed from tract-level US Census data according to the method described in (2). Briefly, participant addresses were linked to tract-level data from the US Census Bureau American Community Survey averaged over five years (https://www.census.gov/programs-surveys/acs). A composite score of neighbourhood-level socioeconomic disadvantage was computed from tract-level proportions of residents reported as unemployed, living below the federal poverty threshold, having less than 12 years of education, not being employed in a management position, and single mothers. These were aggregated to form a neighbourhood-level socioeconomic disadvantage composite measure described in (2), and coded such that higher scores reflect greater disadvantage.

##### Neighborhood opportunity

The neighborhood opportunity measure indexed the intergenerational economic mobility of children of low-income parents. It examines average annual household income in 2014-15 of offspring (born between 1978-1983, who are now in their mid-thirties) of low-income parents (defined as mean pre-tax income at the household level across five years (1994, 1995, 1998-2000) at the 25th percentile of the national income distribution, or $27000/year) within each census tract. Household income was obtained from federal tax return records between 1989-2015, the 2000 and 2010 Decennial Census (US Census Bureau, 2000, 2010; https://data2.nhgis.org/main), and 2005-2015 American Community Surveys (https://www.census.gov/programs-surveys/acs). Census tracts reflect where the child resided through the age of 23. This data was compiled by and obtained from the Opportunity Atlas (https://opportunityatlas.org; 53).

#### Developmental covariates

##### Body mass index (BMI)

BMI is socially-patterned with high BMI being more common in children from lower socioeconomic status families and neighborhoods (56). BMI is also associated with differential DNA-methylation patterns across the genome (57). We therefore considered BMI in our analysis. We measured BMI from in-laboratory measurements of height and weight transformed to gender- and age-normed z-scores according to the method published by the US Centers for Disease Control and Prevention (https://www.cdc.gov/growthcharts/percentile_data_files.htm).

##### Pubertal developmen

Puberty is sometimes reported to onset at younger ages in children growing up in conditions of socioeconomic disadvantage (58). Puberty is also associated with a range of DNA-methylation changes (59, 60). We therefore consider children’s pubertal development in our analysis. Pubertal development was measured using children’s self-reports on the Pubertal Development Scale (61). The scale assesses the extent of development across five sex-specific domains (for both: height, body hair growth, skin changes; for girls: onset of menses, breast development; for boys: growth in body hair, deepening of voice). A total pubertal status score will be computed as the average response (1 = “Not yet begun” to 4 = “Has finished changing”) across all items. Pubertal development was residualized for age, gender, and an age by gender interaction.

##### Tobacco exposure

Smoking is a socially-patterned health behavior to which children from lower socioeconomic status families and neighborhoods are disproportionately exposed (62). It is also associated with differential DNA-methylation patterns across the genome (63, 64). We therefore considered tobacco exposure in our analysis. We measured tobacco exposure using a DNA-methylation smoking (DNAm-smoke) score created by summing the product of the weight and the individual beta estimate for each individual at each CpG site significantly associated with smoking in the discovery EWAS (63). Excluding self-reported tobacco users (n=53) did not significantly alter results.

### Statistical analyses

Six cognitive outcomes were examined in each cognitive model: (1) Processing speed, (2) general executive functions, (3) perceptual reasoning, (4) verbal comprehension, (5) reading, and (6) math.

#### Regression models

We performed multilevel, multivariate regression models fit with FIML in *Mplus* 8.2 statistical software (65). To account for nesting of repeated measures within individuals, and multiple twin pairs within families, a sandwich correction was applied to the standard errors in all analyses. All models included a random intercept, representing the family-level intercept of the dependent variable, to correct for non-independence of twins. All models included age, gender, and an age by gender interaction as covariates. We controlled for multiple testing using the Benjamini–Hochberg false discovery rate (FDR) method (66).

## Declarations

### Funding

We gratefully acknowledge all participants of the Texas Twin Project. This research was supported by National Institutes of Health (NIH) grants R01HD083613 and R01HD092548. LR is supported by the German Research Foundation (DFG). BG, KPH, and EMTD are Faculty Research Associates of the Population Research Center at the University of Texas at Austin, which is supported by a NIH grant P2CHD042849. BG and EMTD are members of the Center on Aging and Population Sciences (CAPS) at The University of Texas at Austin, which is supported by NIH grant P30AG066614. KPH and EMTD were also supported by Jacobs Foundation Research Fellowships. LSK is supported by NIH fellowship F32HD105285-01.

### Conflicts of interest

Not applicable.

### Ethics approval

The University of Texas at Austin Institutional Review board granted ethical approval.

### Consent to participate

Informed consent to participate in the study was obtained from all participants and their parent or legal guardian.

### Consent for publication

Not applicable.

### Availability of data and material

Because of the high potential for deductive identification in this special population from a geographically circumscribed area, and the sensitive nature of information collected, data from the Texas Twin Project are not shared with individuals outside of the research team.

### Code availability

Code will be shared by the first author upon request.

### Author Contributions

KPH, EMTD, & LR developed the study concept and design. LR, PT, AS, TM & LV performed the data analysis under the supervision of EMTD and KPH. LR drafted the manuscript. All authors provided critical revisions and approved the final version of the manuscript for submission.

## Supplement

**Table S1.** Associations between socioeconomic inequality and racial/ethnic identity with DNA-methylation profiles.

|  | DNAm-CRP |  |  | Epigenetic-g |  |  | DunedinPoAm |  |  |
| --- | --- | --- | --- | --- | --- | --- | --- | --- | --- |
|  | *b* | 95% CI | *p* | *b* | 95% CI | *p* | *b* | 95% CI | *p* |
|  | No further covariates <sup>a</sup> |  |  |  |  |  |  |  |  |
| Family-level disadvantage | 0.22 | 0.12– 0.31 | **<0.001** | -0.14 | -0.23 – -0.04 | **0.005** | 0.28 | 0.18– 0.37 | **<0.001** |
| Neighborhood-level disadvantage | 0.25 | 0.17 – 0.33 | **<0.001** | -0.23 | -0.31– -0.15 | **<0.001** | 0.24 | 0.15 – 0.34 | **<0.001** |
| Neighborhood opportunity | -0.12 | -0.21– -0.03 | **0.012** | 0.19 | 0.10 – 0.28 | **<0.001** | -0.19 | -0.30 – -0.08 | **0.001** |
| Latinx | 0.15 | 0.07– 0.24 | **0.001** | -0.10 | -0.17– -0.02 | **0.012** | 0.18 | 0.10– 0.27 | **<0.001** |
| Latinx-White | 0.10 | -0.02– 0.22 | 0.108 | -0.08 | -0.17 – 0.01 | 0.081 | 0.09 | -0.01– 0.19 | 0.089 |
| Black+ | 0.08 | 0.02 – 0.15 | **0.012** | -0.30 | -0.38– -0.22 | **<0.001** | 0.19 | 0.11– 0.28 | **<0.001** |
| Asian+ | -0.01 | -0.09– 0.07 | 0.776 | -0.10 | -0.17– -0.03 | **0.004** | 0.11 | -0.01– 0.22 | 0.058 |
|  | Controlling for BMI |  |  |  |  |  |  |  |  |
| BMI | 0.33 | 0.24– 0.42 | **<0.001** | -0.13 | -0.22– -0.05 | **0.001** | 0.30 | 0.21– 0.40 | **<0.001** |
| Family-level disadvantage | 0.11 | 0.01– 0.21 | 0.024 | -0.08 | -0.17 – 0.01 | 0.076 | 0.19 | 0.08 – 0.29 | **<0.001** |
| Neighborhood-level disadvantage | 0.15 | 0.07– 0.23 | **<0.001** | -0.20 | -0.27– -0.12 | **<0.001** | 0.14 | 0.05 – 0.24 | **0.004** |
| Neighborhood opportunity | -0.05 | -0.15– 0.05 | 0.314 | 0.16 | 0.07 – 0.25 | **<0.001** | -0.09 | -0.20 – 0.01 | 0.088 |
| Latinx | 0.10 | 0.01– 0.18 | 0.029 | -0.08 | -0.16– -0.01 | **0.027** | 0.13 | 0.04– 0.22 | **0.004** |
| Latinx-White | 0.07 | -0.03– 0.17 | 0.180 | -0.07 | -0.15 – 0.02 | 0.150 | 0.06 | -0.03 – 0.14 | 0.213 |
| Black+ | 0.01 | -0.07– 0.07 | 0.975 | -0.25 | -0.33 – -0.17 | **<0.001** | 0.11 | 0.02 – 0.19 | **0.011** |
| Asian+ | 0.01 | -0.07– 0.09 | 0.857 | -0.10 | -0.17 – -0.04 | **0.003** | 0.13 | 0.02 – 0.24 | **0.025** |
|  | Controlling for puberty |  |  |  |  |  |  |  |  |
| Puberty | 0.13 | 0.02– 0.24 | **0.020** | 0.04 | -0.05– 0.13 | 0.392 | 0.05 | -0.07– 0.18 | 0.402 |
| Family-level disadvantage | 0.21 | 0.12 – 0.31 | **<0.001** | -0.12 | -0.22– -0.03 | **0.011** | 0.28 | 0.18– 0.39 | **<0.001** |
| Neighborhood-level disadvantage | 0.25 | 0.16– 0.34 | **<0.001** | -0.23 | -0.31 – -0.15 | **<0.001** | 0.25 | 0.11– 0.34 | **<0.001** |
| Neighborhood opportunity | -0.12 | -0.22 – -0.01 | **0.027** | 0.20 | 0.11– 0.30 | **<0.001** | -0.17 | -0.28– -0.06 | **0.003** |
| Latinx | 0.15 | 0.06– 0.24 | **0.001** | -0.10 | -0.18– -0.03 | **0.008** | 0.18 | 0.09– 0.27 | **<0.001** |
| Latinx-White | 0.14 | 0.01 – 0.26 | **0.033** | -0.08 | -0.17– 0.01 | 0.088 | 0.11 | 0.01– 0.21 | **0.035** |
| Black+ | 0.09 | 0.03 – 0.16 | **0.007** | -0.28 | -0.36– -0.20 | **<0.001** | 0.19 | 0.11– 0.27 | **<0.001** |
| Asian+ | -0.01 | -0.09 – 0.07 | 0.801 | -0.10 | -0.17– -0.03 | **0.006** | 0.11 | -0.01 – 0.22 | 0.059 |
|  | Controlling for DNAm-smoke |  |  |  |  |  |  |  |  |
| DNAm-smoke | 0.40 | 0.28– 0.51 | **<0.001** | 0.34 | 0.22– 0.45 | **<0.001** | 0.24 | 0.11 – 0.37 | **<0.001** |
| Family-level disadvantage | 0.21 | 0.11– 0.30 | **<0.001** | -0.11 | -0.21 – -0.02 | **0.017** | 0.28 | 0.18– 0.38 | **<0.001** |
| Neighborhood-level disadvantage | 0.22 | 0.14– 0.30 | **<0.001** | -0.24 | -0.31– -0.16 | **<0.001** | 0.23 | 0.14 – 0.33 | **<0.001** |
| Neighborhood opportunity | -0.11 | -0.21– -0.02 | **0.022** | 0.19 | 0.10– 0.27 | **<0.001** | -0.17 | -0.27– -0.06 | **0.003** |
| Latinx | 0.11 | 0.03 – 0.20 | **0.011** | -0.13 | -0.20– -0.05 | **0.001** | 0.16 | 0.07– 0.25 | **0.001** |
| Latinx-White | 0.12 | 0.01 – 0.23 | 0.048 | -0.07 | -0.16– 0.01 | 0.093 | 0.11 | 0.01 – 0.20 | **0.028** |
| Black+ | 0.14 | 0.07 – 0.20 | **<0.001** | -0.23 | -0.31– -0.14 | **<0.001** | 0.22 | 0.14 – 0.31 | **<0.001** |
| Asian+ | -0.02 | -0.09 – 0.06 | 0.677 | -0.10 | -0.16– -0.03 | **0.004** | 0.11 | -0.01 – 0.22 | 0.067 |
| Controlling for family-level disadvantage |  |  |  |  |  |  |  |  |  |
|  | *b* | 95% CI | *p* | *b* | 95% CI | *p* | *b* | 95% CI | *p* |
| Family-level disadvantage | 0.19 | 0.07– 0.30 | **0.002** | -0.01 | -0.11– 0.08 | 0.764 | 0.24 | 0.13 – 0.36 | **<0.001** |
| Latinx | 0.10 | 0.01– 0.20 | 0.036 | -0.10 | -0.17– -0.02 | **0.018** | 0.12 | 0.02 – 0.22 | **0.019** |
| Latinx-White | 0.12 | 0.00 – 0.23 | 0.050 | -0.08 | -0.17– 0.01 | 0.081 | 0.10 | 0.01 – 0.20 | 0.047 |
| Black+ | 0.01 | -0.06 – 0.09 | 0.739 | -0.27 | -0.35– -0.19 | **<0.001** | 0.09 | -0.01 – 0.18 | 0.057 |
| Asian+ | 0.01 | -0.08 – 0.09 | 0.914 | -0.10 | -0.17– -0.03 | **0.005** | 0.13 | 0.02 – 0.24 | **0.021** |

Standardized regression coefficients (*b*), 95% confidence intervals (CIs), and uncorrected *p*-values calculated by regressing DNA-methylation measures on socioeconomic measures and racial/ethnic identity with and without controlling for normed BMI z-scores, puberty (residualized for age within each gender), and DNA-methylation profiles of smoking (DNAm-smoke), separately. *P*-values, where FDR corrected *p*-values < 0.05, are marked in bold. <sup>a</sup>All models included covariate adjustment for child's age, gender, and an age by gender interaction. Methylation scores were residualized for technical covariates (array, slide, batch, cell composition).

**Table S2.** Associations between racial/ethnic identity and dimensions of socioeconomic inequality.

|  | Family-level disadvantage |  |  | Neighborhood-level disadvantage |  |  | Neighborhood opportunity |  |  |
| --- | --- | --- | --- | --- | --- | --- | --- | --- | --- |
|  | *b* | 95% CI | *p* | *b* | 95% CI | *p* | *b* | 95% CI | *p* |
| Latinx | 0.25 | 0.16 – 0.35 | **<0.001** | 0.43 | 0.33– 0.52 | **<0.001** | -0.29 | -0.41– -0.17 | **<0.001** |
| Latinx-White | 0.02 | -0.09 – 0.12 | 0.763 | 0.04 | -0.06– 0.15 | 0.444 | 0.03 | -0.11– 0.16 | 0.700 |
| Black+ | 0.43 | 0.35 – 0.52 | **<0.001** | 0.33 | 0.25– 0.41 | **<0.001** | -0.28 | -0.38– -0.18 | **<0.001** |
| Asian+ | -0.10 | -0.19– -0.02 | **0.017** | -0.06 | -0.13– -0.01 | **0.037** | 0.09 | 0.01– 0.17 | 0.040 |

Standardized regression coefficients (*b*), 95% confidence intervals (CIs), and uncorrected *p*-values calculated by regressing socioeconomic measures on racial/ethnic identity. Participants self-identified as White only (62%), Latinx only (12.2%), Latinx-White (8.1%), Black and potentially another race/ethnicity (10%), Asian and potentially another race/ethnicity but not Latinx or Black (7.5%), and Indigenous American, Pacific Islander or other, but not Latinx, Black, or Asian (0.6%, part of reference group due to small sample size). White-only identity is reference group. *P*-values, where FDR corrected *p*-values < 0.05, are marked in bold.

**Table S3.**
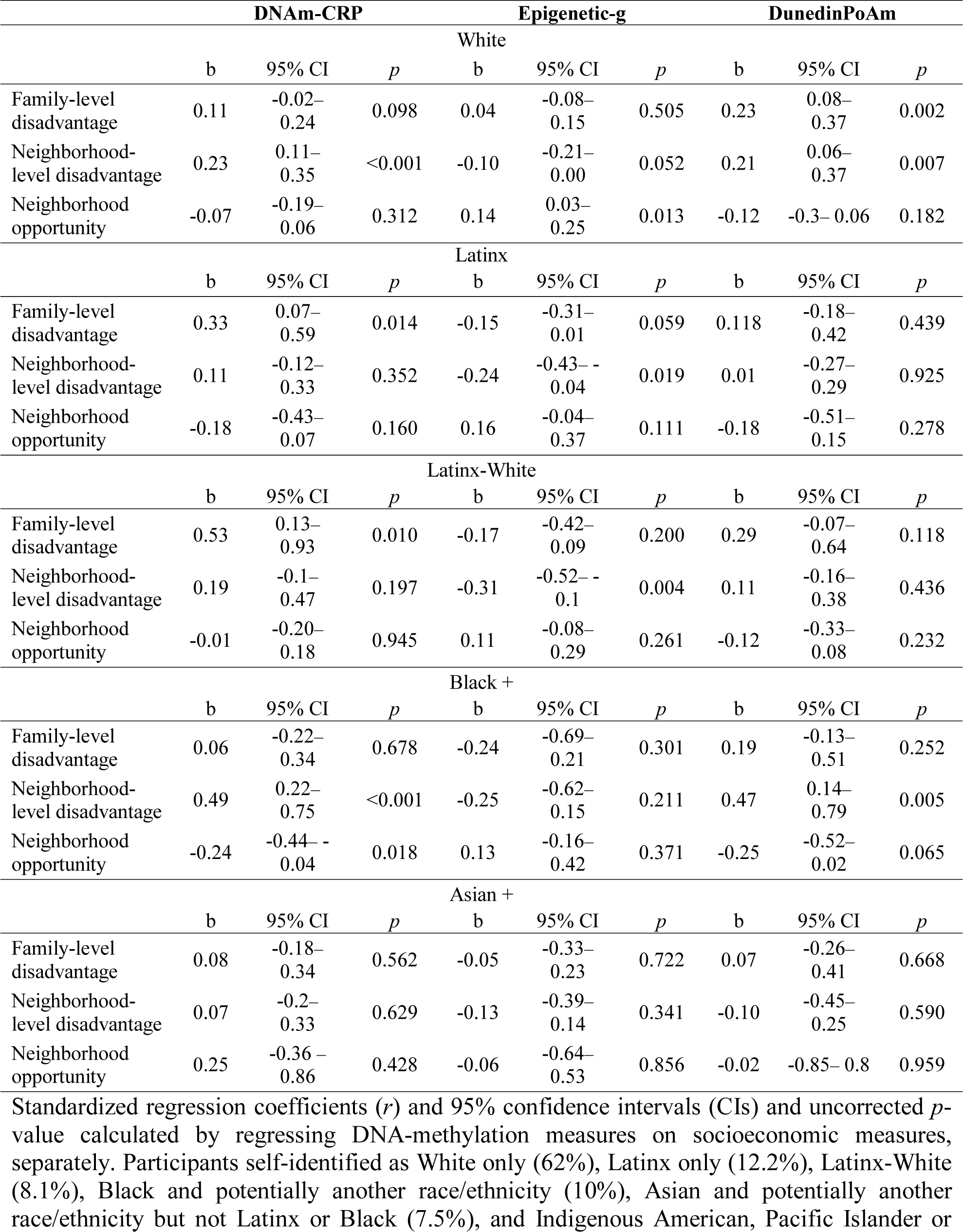
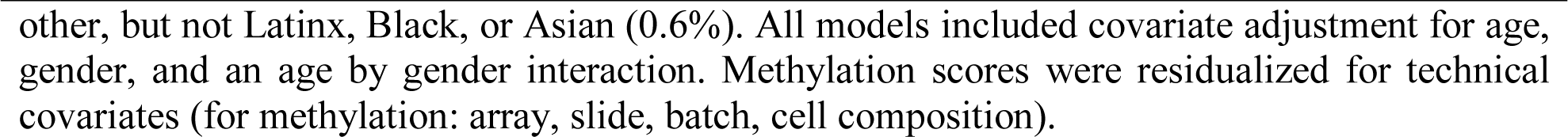
Associations between socioeconomic inequality with DNA-methylation profiles for each racial/ethnic group.

**Table S4.**
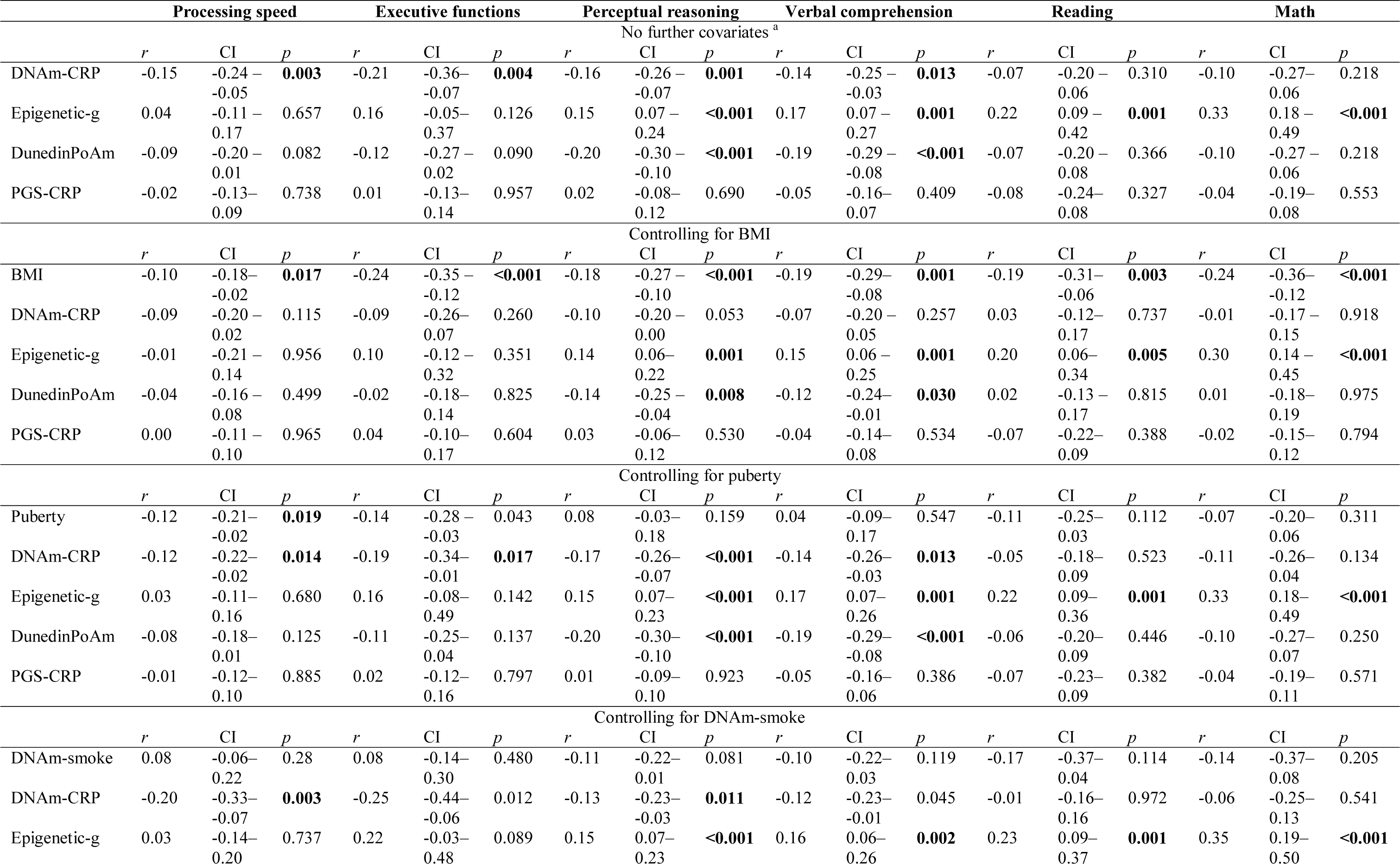
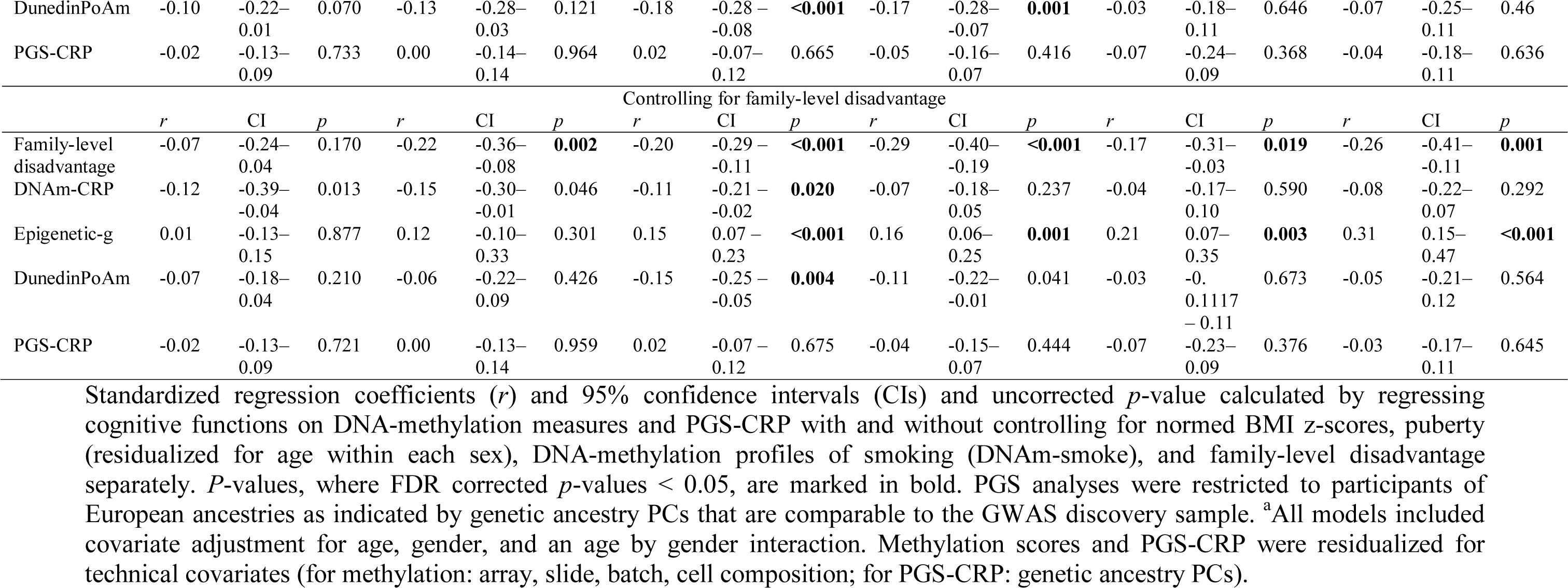
Associations between DNA-methylation and genetic profiles with cognitive functions.

|  | Processing speed |  |  | Executive functions |  |  | Perceptual reasoning |  |  | Verbal comprehension |  |  | Reading |  |  | Math |  |  |
| --- | --- | --- | --- | --- | --- | --- | --- | --- | --- | --- | --- | --- | --- | --- | --- | --- | --- | --- |
|  | No further covariates <sup>a</sup> |  |  |  |  |  |  |  |  |  |  |  |  |  |  |  |  |  |
|  | *r* | CI | *p* | *r* | CI | *p* | *r* | CI | *p* | *r* | CI | *p* | *r* | CI | *p* | *r* | CI | *p* |
| DNA<sub>m</sub>-CRP | -0.15 | -0.24 –<br>-0.05 | **0.003** | -0.21 | -0.36 –<br>-0.07 | **0.004** | -0.16 | -0.26 –<br>-0.07 | **0.001** | -0.14 | -0.25 –<br>-0.03 | **0.013** | -0.07 | -0.20 –<br>0.06 | 0.310 | -0.10 | -0.27 –<br>0.06 | 0.218 |
| Epigenetic-g | 0.04 | -0.11 –<br>0.17 | 0.657 | 0.16 | -0.05 –<br>0.37 | 0.126 | 0.15 | 0.07 –<br>0.24 | **<0.001** | 0.17 | 0.07 –<br>0.27 | **0.001** | 0.22 | 0.09 –<br>0.42 | **0.001** | 0.33 | 0.18 –<br>0.49 | **<0.001** |
| DunedinPoAm | -0.09 | -0.20 –<br>0.01 | 0.082 | -0.12 | -0.27 –<br>0.02 | 0.090 | -0.20 | -0.30 –<br>-0.10 | **<0.001** | -0.19 | -0.29 –<br>-0.08 | **<0.001** | -0.07 | -0.20 –<br>0.08 | 0.366 | -0.10 | -0.27 –<br>0.06 | 0.218 |
| PGS-CRP | -0.02 | -0.13 –<br>0.09 | 0.738 | 0.01 | -0.13 –<br>0.14 | 0.957 | 0.02 | -0.08 –<br>0.12 | 0.690 | -0.05 | -0.16 –<br>0.07 | 0.409 | -0.08 | -0.24 –<br>0.08 | 0.327 | -0.04 | -0.19 –<br>0.08 | 0.553 |
|  | Controlling for BMI |  |  |  |  |  |  |  |  |  |  |  |  |  |  |  |  |  |
|  | *r* | CI | *p* | *r* | CI | *p* | *r* | CI | *p* | *r* | CI | *p* | *r* | CI | *p* | *r* | CI | *p* |
| BMI | -0.10 | -0.18 –<br>-0.02 | **0.017** | -0.24 | -0.35 –<br>-0.12 | **<0.001** | -0.18 | -0.27 –<br>-0.10 | **<0.001** | -0.19 | -0.29 –<br>-0.08 | **0.001** | -0.19 | -0.31 –<br>-0.06 | **0.003** | -0.24 | -0.36 –<br>-0.12 | **<0.001** |
| DNA<sub>m</sub>-CRP | -0.09 | -0.20 –<br>0.02 | 0.115 | -0.09 | -0.26 –<br>0.07 | 0.260 | -0.10 | -0.20 –<br>0.00 | 0.053 | -0.07 | -0.20 –<br>0.05 | 0.257 | 0.03 | -0.12 –<br>0.17 | 0.737 | -0.01 | -0.17 –<br>0.15 | 0.918 |
| Epigenetic-g | -0.01 | -0.21 –<br>0.14 | 0.956 | 0.10 | -0.12 –<br>0.32 | 0.351 | 0.14 | 0.06 –<br>0.22 | **0.001** | 0.15 | 0.06 –<br>0.25 | **0.001** | 0.20 | 0.06 –<br>0.34 | **0.005** | 0.30 | 0.14 –<br>0.45 | **<0.001** |
| DunedinPoAm | -0.04 | -0.16 –<br>0.08 | 0.499 | -0.02 | -0.18 –<br>0.14 | 0.825 | -0.14 | -0.25 –<br>-0.04 | **0.008** | -0.12 | -0.24 –<br>-0.01 | **0.030** | 0.02 | -0.13 –<br>0.17 | 0.815 | 0.01 | -0.18 –<br>0.19 | 0.975 |
| PGS-CRP | 0.00 | -0.11 –<br>0.10 | 0.965 | 0.04 | -0.10 –<br>0.17 | 0.604 | 0.03 | -0.06 –<br>0.12 | 0.530 | -0.04 | -0.14 –<br>0.08 | 0.534 | -0.07 | -0.22 –<br>0.09 | 0.388 | -0.02 | -0.15 –<br>0.12 | 0.794 |
|  | Controlling for puberty |  |  |  |  |  |  |  |  |  |  |  |  |  |  |  |  |  |
|  | *r* | CI | *p* | *r* | CI | *p* | *r* | CI | *p* | *r* | CI | *p* | *r* | CI | *p* | *r* | CI | *p* |
| Puberty | -0.12 | -0.21 –<br>-0.02 | **0.019** | -0.14 | -0.28 –<br>-0.03 | 0.043 | 0.08 | -0.03 –<br>0.18 | 0.159 | 0.04 | -0.09 –<br>0.17 | 0.547 | -0.11 | -0.25 –<br>0.03 | 0.112 | -0.07 | -0.20 –<br>0.06 | 0.311 |
| DNA<sub>m</sub>-CRP | -0.12 | -0.22 –<br>-0.02 | **0.014** | -0.19 | -0.34 –<br>-0.01 | **0.017** | -0.17 | -0.26 –<br>-0.07 | **<0.001** | -0.14 | -0.26 –<br>-0.03 | **0.013** | -0.05 | -0.18 –<br>0.09 | 0.523 | -0.11 | -0.26 –<br>0.04 | 0.134 |
| Epigenetic-g | 0.03 | -0.11 –<br>0.16 | 0.680 | 0.16 | -0.08 –<br>0.49 | 0.142 | 0.15 | 0.07 –<br>0.23 | **<0.001** | 0.17 | 0.07 –<br>0.26 | **0.001** | 0.22 | 0.09 –<br>0.36 | **0.001** | 0.33 | 0.18 –<br>0.49 | **<0.001** |
| DunedinPoAm | -0.08 | -0.18 –<br>0.01 | 0.125 | -0.11 | -0.25 –<br>0.04 | 0.137 | -0.20 | -0.30 –<br>-0.10 | **<0.001** | -0.19 | -0.29 –<br>-0.08 | **<0.001** | -0.06 | -0.20 –<br>0.09 | 0.446 | -0.10 | -0.27 –<br>0.07 | 0.250 |
| PGS-CRP | -0.01 | -0.12 –<br>0.10 | 0.885 | 0.02 | -0.12 –<br>0.16 | 0.797 | 0.01 | -0.09 –<br>0.10 | 0.923 | -0.05 | -0.16 –<br>0.06 | 0.386 | -0.07 | -0.23 –<br>0.09 | 0.382 | -0.04 | -0.19 –<br>0.11 | 0.571 |
|  | Controlling for DNAm-smoke |  |  |  |  |  |  |  |  |  |  |  |  |  |  |  |  |  |
|  | *r* | CI | *p* | *r* | CI | *p* | *r* | CI | *p* | *r* | CI | *p* | *r* | CI | *p* | *r* | CI | *p* |
| DNAm-smoke | 0.08 | -0.06 –<br>0.22 | 0.28 | 0.08 | -0.14 –<br>0.30 | 0.480 | -0.11 | -0.22 –<br>0.01 | 0.081 | -0.10 | -0.22 –<br>0.03 | 0.119 | -0.17 | -0.37 –<br>0.04 | 0.114 | -0.14 | -0.37 –<br>0.08 | 0.205 |
| DNAm-CRP | -0.20 | -0.33 –<br>-0.07 | **0.003** | -0.25 | -0.44 –<br>-0.06 | 0.012 | -0.13 | -0.23 –<br>-0.03 | **0.011** | -0.12 | -0.23 –<br>-0.01 | 0.045 | -0.01 | -0.16 –<br>0.16 | 0.972 | -0.06 | -0.25 –<br>0.13 | 0.541 |
| Epigenetic-g | 0.03 | -0.14 –<br>0.20 | 0.737 | 0.22 | -0.03 –<br>0.48 | 0.089 | 0.15 | 0.07 –<br>0.23 | **<0.001** | 0.16 | 0.06 –<br>0.26 | **0.002** | 0.23 | 0.09 –<br>0.37 | **0.001** | 0.35 | 0.19 –<br>0.50 | **<0.001** |
| DunedinPoAm | -0.10 | -0.22–<br>0.01 | 0.070 | -0.13 | -0.28–<br>0.03 | 0.121 | -0.18 | -0.28 –<br>-0.08 | **<0.001** | -0.17 | -0.28–<br>-0.07 | **0.001** | -0.03 | -0.18–<br>0.11 | 0.646 | -0.07 | -0.25–<br>0.11 | 0.46 |
| PGS-CRP | -0.02 | -0.13–<br>0.09 | 0.733 | 0.00 | -0.14–<br>0.14 | 0.964 | 0.02 | -0.07–<br>0.12 | 0.665 | -0.05 | -0.16–<br>0.07 | 0.416 | -0.07 | -0.24–<br>0.09 | 0.368 | -0.04 | -0.18–<br>0.11 | 0.636 |
| Controlling for family-level disadvantage |  |  |  |  |  |  |  |  |  |  |  |  |  |  |  |  |  |  |
|  | *r* | CI | *p* | *r* | CI | *p* | *r* | CI | *p* | *r* | CI | *p* | *r* | CI | *p* | *r* | CI | *p* |
| Family-level disadvantage | -0.07 | -0.24–<br>0.04 | 0.170 | -0.22 | -0.36–<br>-0.08 | **0.002** | -0.20 | -0.29 –<br>-0.11 | **<0.001** | -0.29 | -0.40–<br>-0.19 | **<0.001** | -0.17 | -0.31–<br>-0.03 | **0.019** | -0.26 | -0.41–<br>-0.11 | **0.001** |
| DNA-m-CRP | -0.12 | -0.39–<br>-0.04 | 0.013 | -0.15 | -0.30–<br>-0.01 | 0.046 | -0.11 | -0.21 –<br>-0.02 | **0.020** | -0.07 | -0.18–<br>0.05 | 0.237 | -0.04 | -0.17–<br>0.10 | 0.590 | -0.08 | -0.22–<br>0.07 | 0.292 |
| Epigenetic-g | 0.01 | -0.13–<br>0.15 | 0.877 | 0.12 | -0.10–<br>0.33 | 0.301 | 0.15 | 0.07 –<br>0.23 | **<0.001** | 0.16 | 0.06–<br>0.25 | **0.001** | 0.21 | 0.07–<br>0.35 | **0.003** | 0.31 | 0.15–<br>0.47 | **<0.001** |
| DunedinPoAm | -0.07 | -0.18–<br>0.04 | 0.210 | -0.06 | -0.22–<br>0.09 | 0.426 | -0.15 | -0.25 –<br>-0.05 | **0.004** | -0.11 | -0.22–<br>-0.01 | 0.041 | -0.03 | -0.11–<br>0.11 | 0.673 | -0.05 | -0.21–<br>0.12 | 0.564 |
| PGS-CRP | -0.02 | -0.13–<br>0.09 | 0.721 | 0.00 | -0.13–<br>0.14 | 0.959 | 0.02 | -0.07 –<br>0.12 | 0.675 | -0.04 | -0.15–<br>0.07 | 0.444 | -0.07 | -0.23–<br>0.09 | 0.376 | -0.03 | -0.17–<br>0.11 | 0.645 |

Standardized regression coefficients (*r*) and 95% confidence intervals (CIs) and uncorrected *p*-value calculated by regressing cognitive functions on DNA-methylation measures and PGS-CRP with and without controlling for normed BMI z-scores, puberty (residualized for age within each sex), DNA-methylation profiles of smoking (DNA-m-smoke), and family-level disadvantage separately. *P*-values, where FDR corrected *p*-values < 0.05, are marked in bold. PGS analyses were restricted to participants of European ancestries as indicated by genetic ancestry PCs that are comparable to the GWAS discovery sample. <sup>a</sup>All models included covariate adjustment for age, gender, and an age by gender interaction. Methylation scores and PGS-CRP were residualized for technical covariates (for methylation: array, slide, batch, cell composition; for PGS-CRP: genetic ancestry PCs).

**Table S5.**
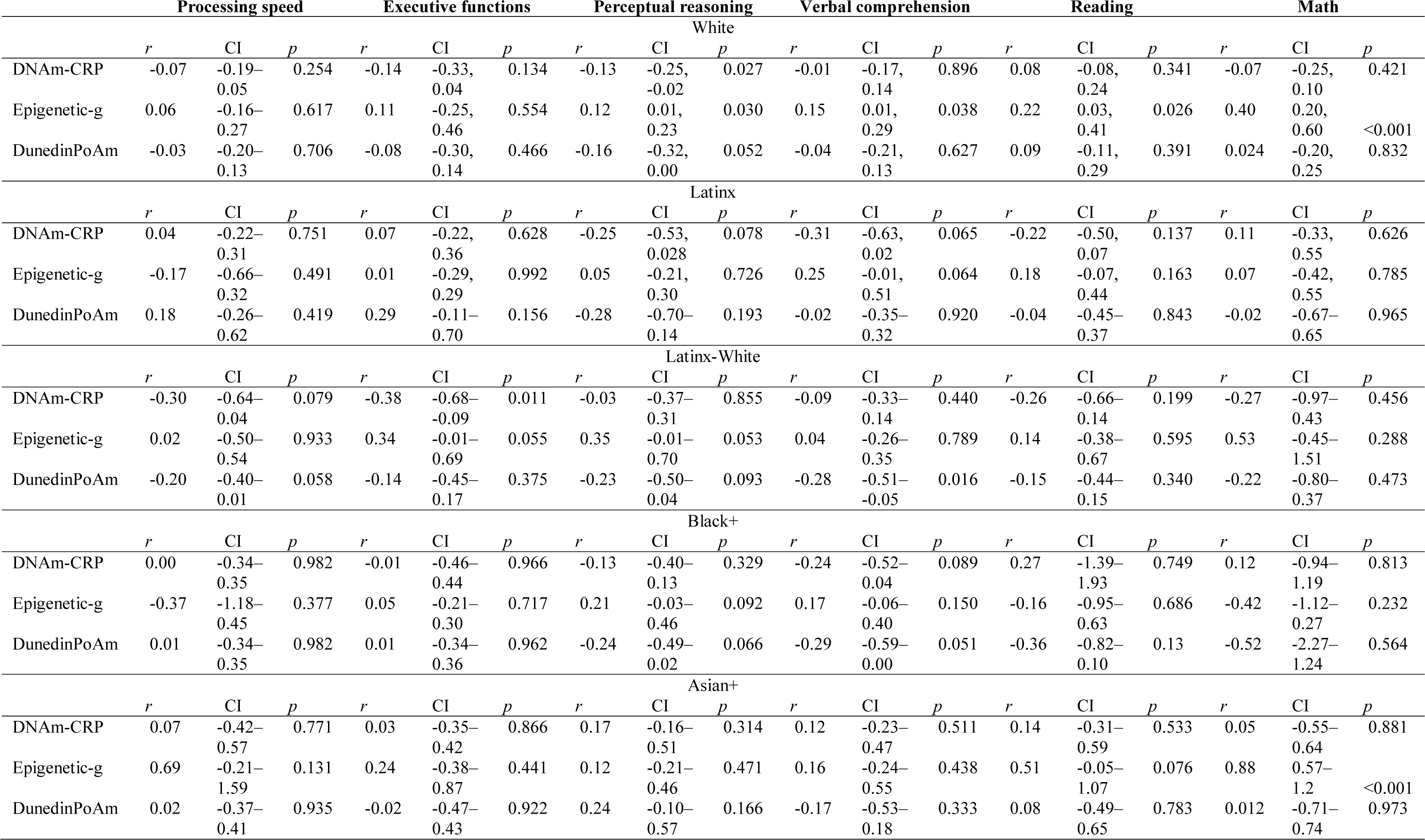
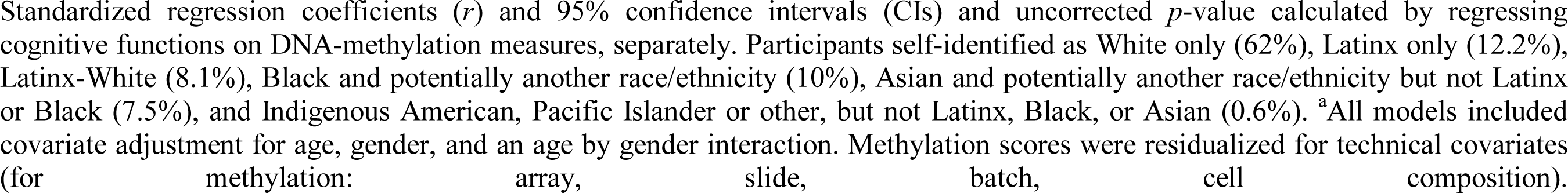
Associations between DNA-methylation with cognitive functions for each racial/ethnic group.

**Table S6.**
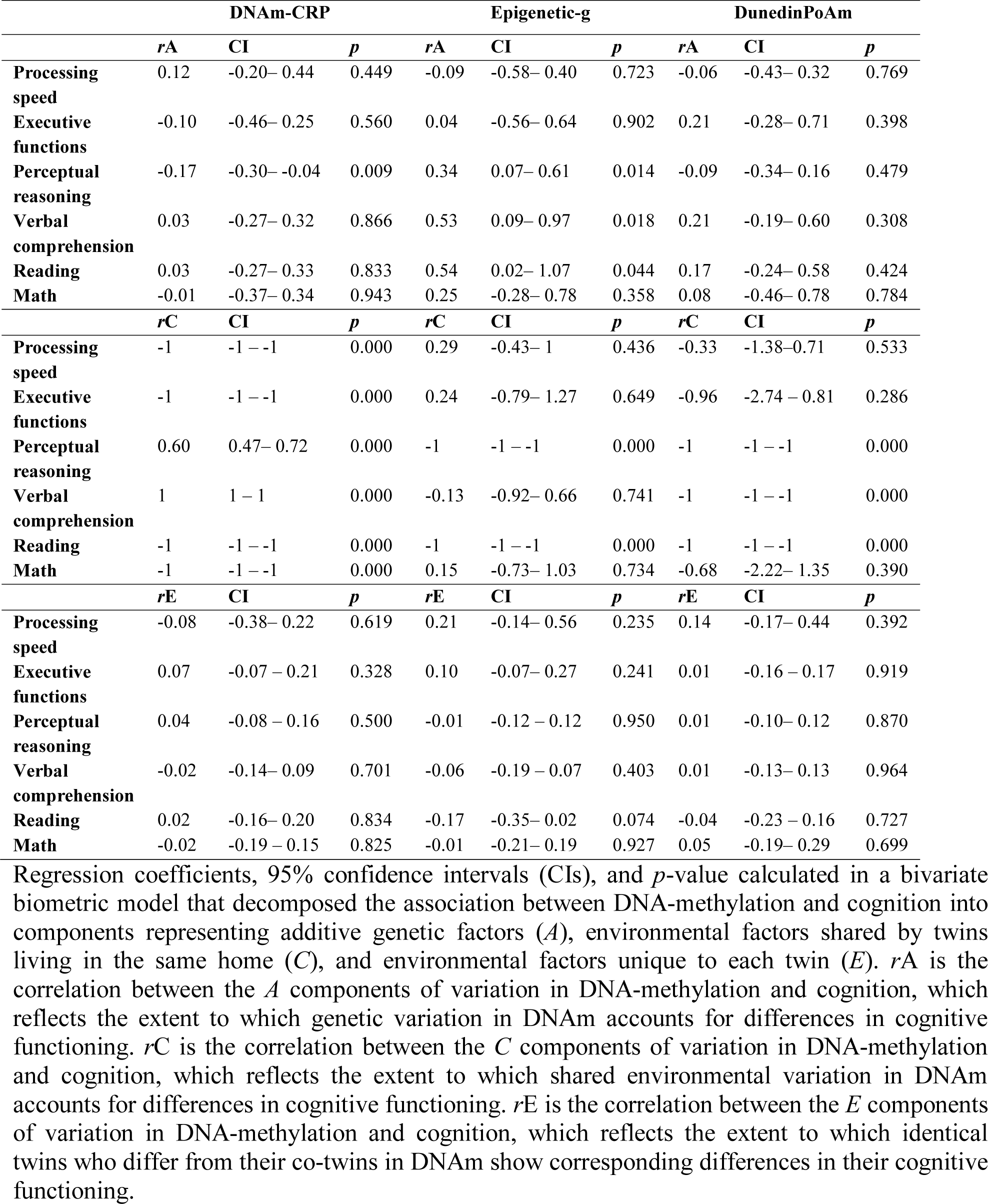
Co-twin-control associations between DNA-methylation and cognitive functions.

**Figure S1.**
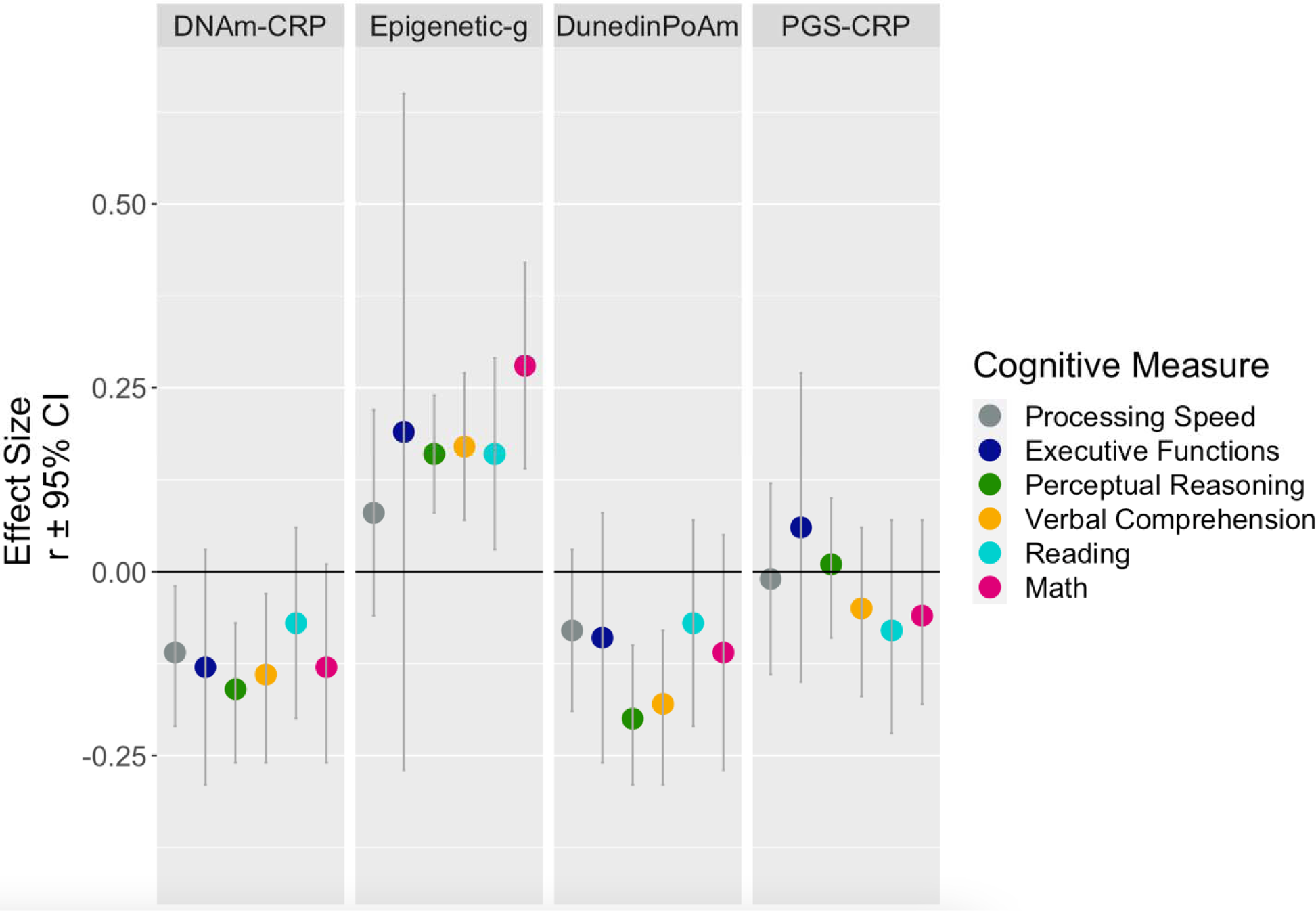
Associations between three DNA-methylation profiles (DNAm-CRP, Epigenetic-g, and DunedinPoAm) and a polygenic score of inflammation (PGS-CRP) with six measures of cognitive functioning in children and adolescents with listwise deletion. The plot depicts the standardized regression coefficients (*r*) and 95% confidence intervals (CIs) calculated by regressing cognitive functions on DNA-methylation measures and PGS-CRP, separately with listwise deletion. PGS analyses were restricted to participants solely of recent European ancestries as indicated by genetic ancestry PCs that are comparable to the GWAS discovery sample. All models included covariate adjustment for child’s age and gender, and technical covariates. Higher cognitive values indicate higher task performance. Higher DNAm-CRP and PGS-CRP values indicate a methylation profile and genetic profile of higher chronic inflammation, respectively. Higher Epigenetic-g values indicate a methylation profile associated with higher cognitive functioning. Higher DunedinPoAm values indicate a methylation profile of faster biological aging.

## Notes

### Competing Interest Statement

The authors have declared no competing interest.

